# Investigating E3 ubiquitin ligase activity of ORF150 in fish herpesvirus pathogenesis

**DOI:** 10.1101/2025.02.21.639420

**Authors:** Hao Wang, Jia Yang, Mengjuan Li, Ye Zhang, Liqun Lu

## Abstract

Fish herpesviruses, members of the family *Alloherpesviridae*, infect aquatic species. The host’s ubiquitin-proteasome system (UPS) serves as the major intracellular pathway for protein degradation and plays a crucial role in the viral replication cycle. Viruses exploit the UPS to facilitate their replication and evade host immune responses. This study demonstrated that cyprinid herpesvirus 2 (CyHV-2) infection significantly activates the host UPS pathway, as revealed by transcriptomics sequencing analysis. Further, treatment with MG132, a proteasome inhibitor, markedly downregulates intracellular viral replication and gene transcription. Additionally, we identified a novel, highly conserved E3 ubiquitin ligase with a C3HC4-type really interesting gene (RING)-finger (RING-HC) domain, encoded by open reading frame 150 (*ORF150*) of CyHV-2. These findings provide critical molecular evidence that the host UPS is essential for fish herpesvirus replication and suggest that viral E3 ligases may serve as potential targets for new antiviral strategies.

## Introduction

Fish herpesviruses present substantial challenges to aquaculture due to their host specificity, environmental persistence, and complex immune interactions (1). Cyprinid herpesvirus 2 (CyHV-2; also known as goldfish hematopoietic necrosis virus), a member of the genus *Cyvirus* of the family *Alloherpesviridae*, along with CyHV-1 (also known as carp pox herpesvirus) and CyHV-3 (also known as koi herpesvirus), has been reported in numerous countries and regions globally (2–4). CyHV-2 induces acute infection within host and exhibits latent properties, further complicating its management by triggering secondary bacterial infections (5–7). The infection mechanism remains complex, and to date, no targeted control strategies have yet been developed against CyHV-2. Although certain compounds and vaccines have shown antiviral efficacy and protective functions in CyHV-2 infections, their development process is lengthy, and their evaluation and practical application timelines remain uncertain (8–10). Until now, the molecular mechanisms underlying CyHV-2 replication are still unclear. Whereas transcriptomic and proteomic analyses have revealed several signaling pathways potentially involved in CyHV-2 infection, including p53, RIG-I-like receptor, JAK/STAT, and PI3K/AKT/mTOR pathway (11, 12), providing valuable clues for further exploration of the CyHV-2 infection mechanism.

The viral genome harbors limited genetic information and relies on host cellular pathways for nearly all stages of its infection cycle (13). The ubiquitin-proteasome system (UPS) serves as the major pathway for intracellular protein degradation, regulating essential cellular processes (14, 15). Specifically, the UPS covalently attaches ubiquitin to target substrates via a sequential cascade involving E1-activating enzymes, E2-conjugating enzymes, and E3 ligases, subsequently, the polyubiquitinated proteins are recognized and degraded by the 26S proteasomes (16). There is increasing evidence that the UPS plays an essential role in the viral replication cycle. In virus-host interactions, both exploit versatile ubiquitination code to compete, combat, and co-evolve (17, 18). Consequently, viruses manipulate and hijack the UPS to benefit their own infectious cycles and evade host immune response (19, 20). For instance, inhibition of proteasome activity reduces Kaposi’s sarcoma-associated herpesvirus (KSHV) entry into endothelial cells and intracellular trafficking (21). Another study confirmed that the UPS is identified as a key role in the life cycle of Singapore grouper iridovirus (SGIV) (22). Moreover, the UPS constitutes a host defense mechanism to eliminate viral replication; in response, viruses may encode their own ubiquitin components, particularly E3 ligases, to counteract this machinery (20, 23). Notably, almost all reported viral E3 ligases belong to the really interesting new gene (RING) family, which might reflect an ascertainment bias (20). Herpes simplex virus 1 (HSV-1) employs its viral E3 ligase infected cell protein 0 (ICP0) to establish its life cycle by modulating host immune responses (24). Similarly, white spot syndrome virus (WSSV), an aquatic viruse, encodes four RING-H2 type E3 ligases (25). Furthermore, whole genome sequencing analysis has revealed potential RING gene families encoding E3 ligases in the genus *Cyvirus*, including CyHVs (26–28). Therefore, investigating the molecular mechanism of interaction between CyHV-2 and the host UPS could provide alternative targets for antiviral strategies.

Herein, we demonstrated that CyHV-2 infection significantly activates the host UPS pathway and the role of proteasomes in viral replication was evaluated utilizing MG132. Furthermore, we identified a novel, highly conserved RING-HC type E3 ligase encoded by open reading frame 150 (*ORF150*) of CyHV-2. This study represents the first molecular evidence that UPS involves in fish herpesvirus infection. These findings provide valuable insights into the functions of viral E3 ligase and the molecular mechanisms underlying fish herpesvirus-host interactions.

## Results

### UPS is activated in CyHV-2 infection

To investigate potential cellular factors involved in CyHV-2 replication, the differentially expressed genes (DEGs) during viral infection were identified through transcriptomics analysis. A total of 107 DEGs related to the UPS components in gibel carp were identified, with 71 upregulated genes and 36 downregulated genes upon CyHV-2 infection (Fig. 1*A*). These genes are involved in ubiquitination, proteasome degradation, and deubiquitination (Fig. 1*B*). Among them, most genes were significantly upregulated, including those encoding the 20S core particles (i.e., *psma1– 7* and *psmb1–7*, which encode the α and β subunits 1–7 of 20S proteasome); the 19S regulatory particles (i.e., *psmc1–6*, which encode the ATPase subunits of the 26S proteasome; *psmd1–4*, *psmd6–8*, and *psmd11b–14*, which encode the non-ATPase regulatory subunits); the proteasome regulator (i.e., *psmf1*, encoding the proteasome inhibitor subunit); the proteasome assembly factors (i.e., *psmg1* and *psmg4*, which encode the proteasome assembly chaperones; *pomp*, encoding the proteasome maturation protein); the proteasome-interacting proteins (i.e., *ecm29*, encoding a proteasome adaptor and scaffold; *psmd5*); deubiquitylating enzymes (DUBs) (e.g., *usp4*, encoding ubiquitin-specific protease 4; *usp14* and *usp18*); ubiquitin-like modifier-activating enzyme 1 (*uba1*); E2 enzymes (e.g., *ube2a*, encoding ubiquitin-conjugating enzyme E2a; *ube2d2* and *ube2n*); and E3 ligases (e.g., *trim6*, encoding tripartite motif containing 6; *rnf20*, encoding RING finger protein 20). In contrast, several genes were significantly downregulated, including those encoding the proteasome regulator (*psme4*); DUBs (e.g., *usp7* and *usp36*); E2 enzyme (i.e., *tmem189*, encoding transmembrane protein 189); and multiple E3 ligases (e.g., *trim2*, *rnf8*, and *rnf130*), as shown in Fig. 1, *B*–*F*. Meanwhile, the expression patterns of selected associated genes was validated in vitro, further corroborating the aforementioned findings (Fig. S1). These results suggest the host UPS is activated during CyHV-2 infection.

**Figure 1.**
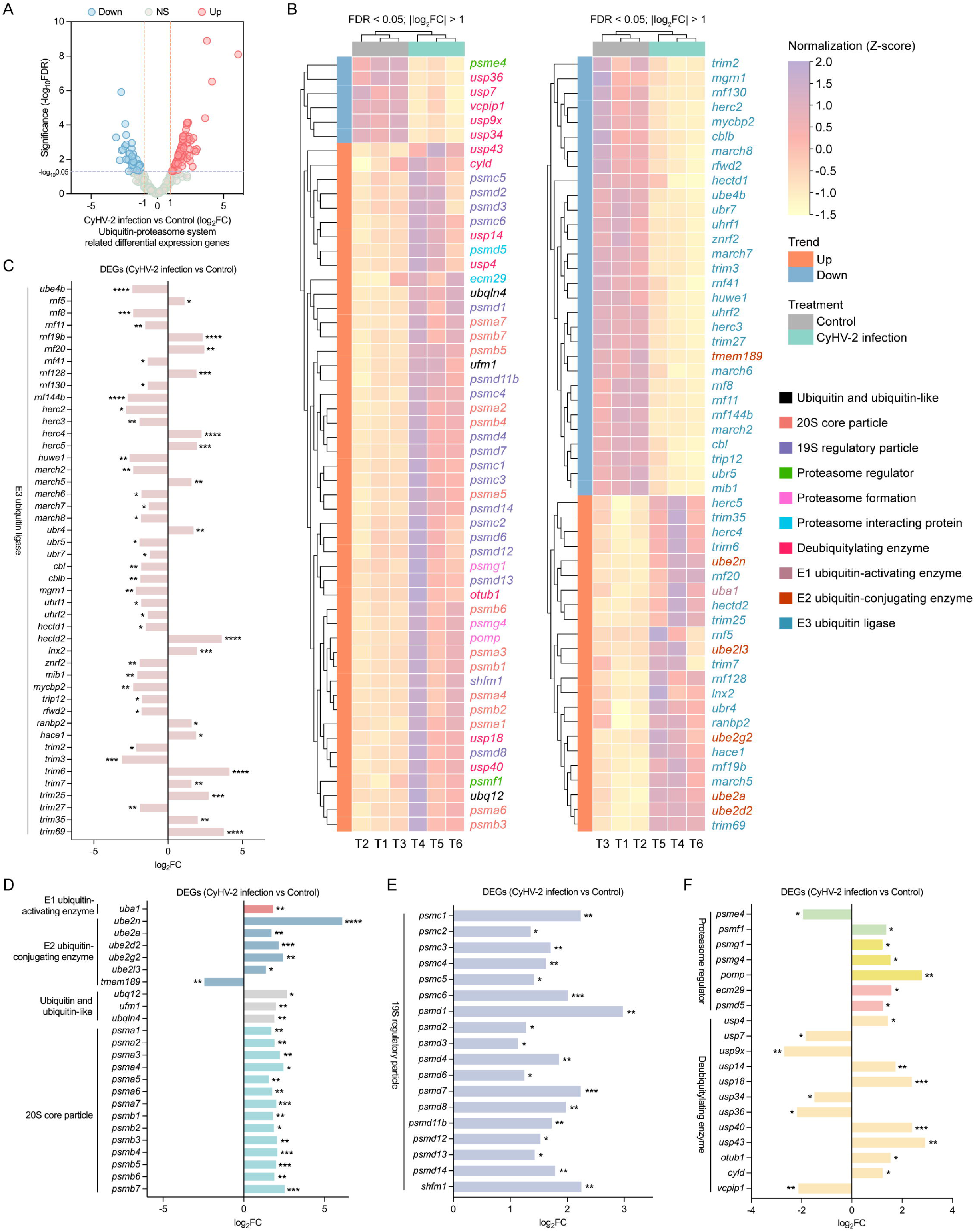
Host ubiquitin-proteasome system (UPS) is activated during cyprinid herpesvirus 2 (CyHV-2) infection through transcriptomics analysis. *A*, volcano plot of UPS-associated differentially expressed genes (DEGs). The x-axis represents log_2_-transformed fold change (log_2_FC; CyHV-2 infection *vs.* control), and the y-axis corresponds to -log_10_ transformed false discover rate (FDR). Vertical orange dashed lines demarcate the threshold for absolute log_2_FC > 1, while the horizontal purple dashed line indicates FDR < 0.05. DEGs are categorized as downregulated (blue), upregulated (red), or non-significant (gray). *B*, hierarchical clustering heatmap of UPS-enriched DEG expression dynamics. Gene expression levels (reads per kilobase of transcript per million mapped reads, RPKM normalized by Z-score and row standardization with a cut-off as description in *A*) are displayed for control (gray) and CyHV-2-infected (cyan) groups, with three biological replicates per condition. Vertical clustering (left) groups genes by expression patterns: upregulated (orange) and downregulated (blue). UPS functional components are color-coded for clarity. In pairwise comparisons between groups, yellow and purple color gradients denote downregulation and upregulation of individual genes, respectively. *C–F*, the log_2_FC values of DEGs are shown in bar graphs with FDR-corrected p-values denoted as * *P* < 0.05, ** *P* < 0.01, *** *P* < 0.001, **** *P* < 0.0001.

### Host proteasome involves in CyHV-2 replication

To further determine whether the UPS is essential for CyHV-2 infection, we employed MG-132 to specifically inhibit proteasome activity. Our results demonstrated that intracellular viral copy numbers were significantly reduced in a dose-dependent manner following MG132 treatment, compared to the dimethyl sulfoxide (DMSO) control (Fig. 2*A*). Furthermore, intracellular viral gene transcriptions were markedly downregulated in a dose-dependent fashion under proteasome inhibition (except for *ORF54*, whose transcription was only significantly decreased with MG132 treatment at 50 μM), affecting several immediate-early (IE), early (E), and late (L) genes, as shown in Fig. 2*B*. Immunofluorescence assay (IFA) observations revealed that viral protein expression was significantly suppressed in the presence of MG132, whereas DMSO-treated cells exhibited widespread viral protein distribution throughout the cell (Fig. 2*C*). These findings confirm that MG132 effectively inhibits intracellular viral transcription and replication, thereby highlighting the critical role of the proteasome in CyHV-2 infection.

**Figure 2.**
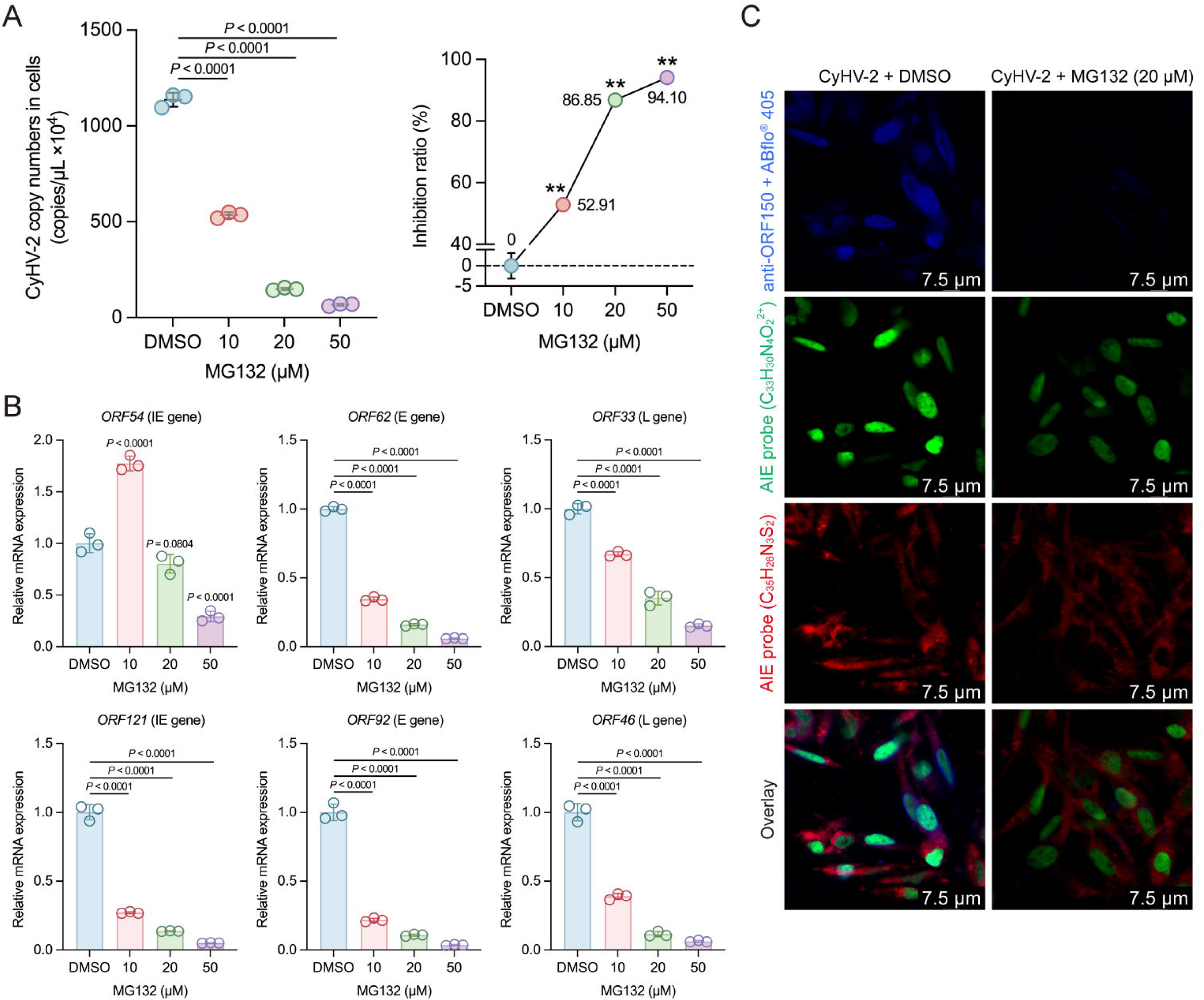
MG132 suppresses intracellular CyHV-2 replication and viral gene transcription. *A* and *B*, dose-dependent antiviral effects of MG132 (10, 20, and 50 μM). Panel *A* (left): viral load quantified by qPCR showing reduced CyHV-2 copy numbers under MG132 treatment; panel *A* (right): inhibition ratios calculated against dimethyl sulfoxide (DMSO) controls (100% viability). Panel *B*: RT-qPCR analysis of viral transcriptional dynamics, including immediate-early (IE), early (E), and late (L) genes. DMSO treatment served as a control. All quantitative data in panels *A* and *B* represent mean ± standard deviation (SD) from three biologically independent experiments. Statistical significance was determined by one-way analysis of variance (ANOVA) shown on the corresponding data in plots or bars. ** *P* < 0.01. *C*, immunofluorescence validation of viral replication suppression. The virions (ORF150 antibody followed by staining with ABflo^®^ 405, cyan) colocalize with nuclei (aggregation-induced emission, i.e., AIE probe, C_33_H_30_N_4_O_2_^2+^, green) and membranes (AIE probe, C_35_H_26_N_3_S_2_, red). Scale bar: 7.5 μm.

### CyHV-2 encodes conserved RING-finger proteins

The host UPS involving in CyHV-2 replication has been confirmed through the aforementioned exploration. We were also interested in the importance of viral components analogous to host UPS homologs. Viruses, particularly those with large DNA genomes such as herpesviruses, may encode their own ubiquitin components, especially E3 ligases, to counteract the host’s antiviral machinery (20, 23). As shown in Fig. 3, *A* and *B*, five RING family genes (*ORF41*, *54*, *128*, *144*, and *150*) have been identified in the CyHV-2 genomes, exhibiting high sequence identities among different isolates and homology within CyHVs (*ORF128* is absent in the SY-C1 strain). This suggests that the RING family is widely present in fish herpesviruses and potentially plays a crucial role in viral infection. As showed in Fig. 3, *C* and *D*, these five proteins all possess a consensus C3HC4 type RING-finger (RING-HC) domain, i.e., CX2CX(9–39)CX(1–3)HX(2–3)C/HX2CX(4–48)CX2C, where cysteine (Cys) and histidine (His) residues serve as zinc-binding sites. The presence of a cysteine residue in the fifth coordination site is indicative of a RING-HC domain, notably, the domain is located at the C-terminus of ORF54 and the N-terminus of other four proteins (Fig. 3*C*). Phylogenetically, these five RING-finger proteins, including ORF150, are homologous and highly conserved across CyHVs and seven CyHV-2 iaolates (Figs. 3*E* and S2*A*). Multiple alignments shown in Figs. 3*F* and S2*B* indicated that these five proteins possess highly consensus RING-HC motifs among CyHVs. These findings suggest that these conserved RING-finger proteins likely play key roles in CyHV-2 infection.

**Figure 3.**
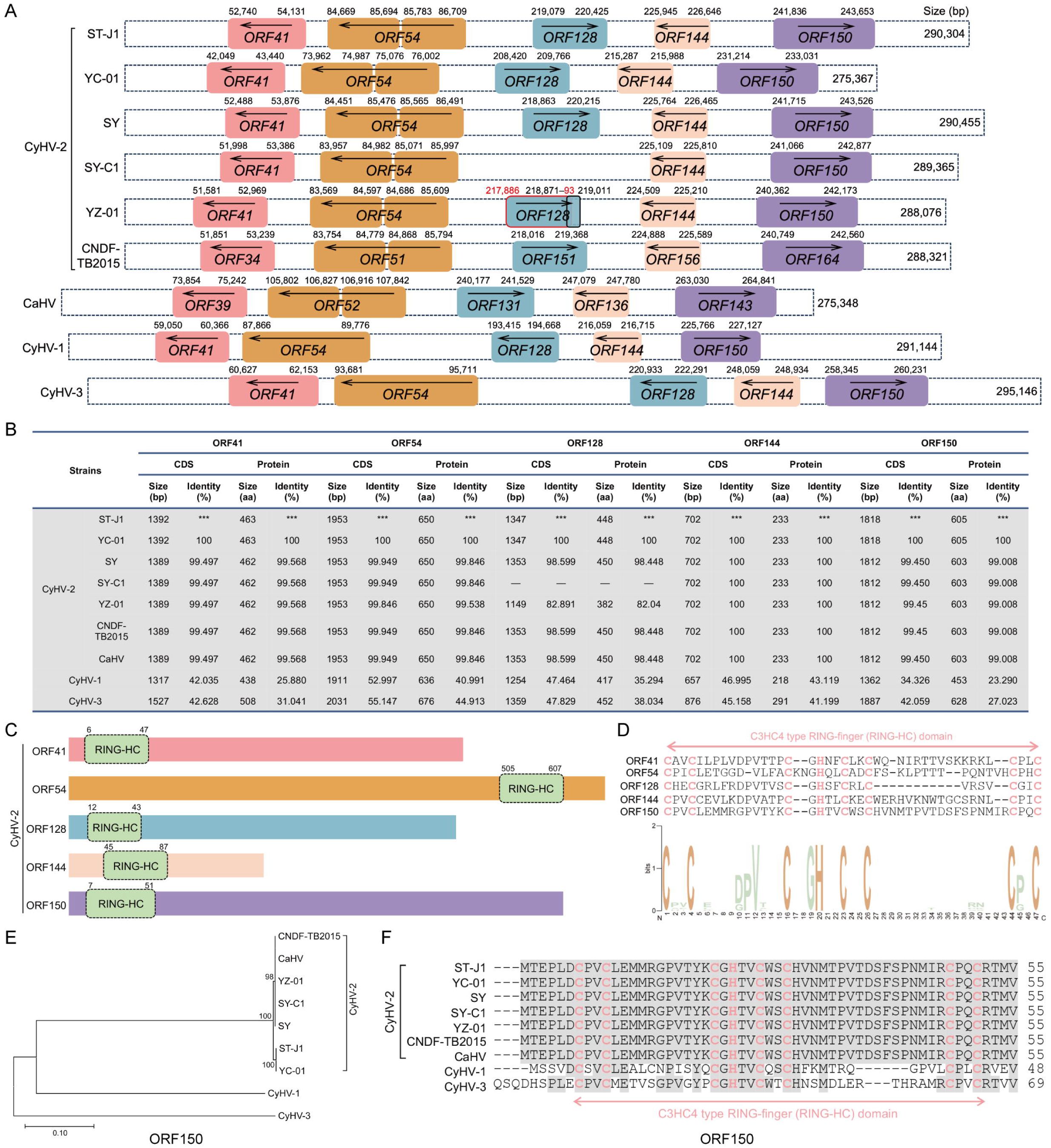
Evolutionary conservation of really interesting new gene (RING)-finger proteins in CyHV-2. *A*, RING-finger family genes layout in CyHVs’ genomes. Dashed-line frames demarcate whole genomes (sizes annotated at frame ends). Colored boxes denote genes encoding RING-finger family members, with directional arrows indicating transcriptional orientation and position coordinates (start/stop codons above boxes). *B*, comparative analysis of RING-finger coding sequences (CDSs): length (bp), translated protein size (aa), and pairwise identity across CyHVs. *C*, domain architecture mapping of five RING-finger proteins in CyHV-2: full-length RING-finger proteins (colored boxes) with embedded RING-HC domains (green sub-boxes, position coordinates above). *D*, the consensus RING-HC domain shown following the alignment of the sequences for the five proteins depicted in panel *C*. The conserved cysteine (C) and histidine (H) amino residues were highlighted in pink. Sequence conservation was visualized by WebLogo 3 (y-axis: bits scale; x-axis: window width), using Shannon entropy-based stacking of amino acid frequencies. *E*, phylogenetic reconstruction of ORF150 homologs across CyHVs. Neighbor-joining tree generated in MEGA v11 (bootstrap = 1000 replicates). *F*, the multiple sequence alignments of ORF150 homologs across CyHVs (complete amino acid sequences are not shown). Grey shading indicates conserved regions and pink boxes mark RING-HC domains. The protein IDs for the ORF150 homologs presented in panels *E* and *F* are listed as follows: CyHV-2 ST-J1, YP_007003966.1; SY-C1, AKC02089.1; SY, AMB21716.1; YC-01, QIV66963.1; YZ-01, QAU54868.1; CNDF-TB2015, QIM55323.1; *Carassius auratus* herpesvirus (CaHV), APB92997.1; CyHV-1, YP_007003805.1; CyHV-3, YP_001096185.1.

### ORF150 functions as a E3 ligase for CyHV-2

We screened and confirmed that the transcriptional expression level of *ORF150* is significantly upregulated during CyHV-2 infection (Fig. S3*A*), suggesting its potential importance for viral replication. Then, we purified ORF150-His (approximately 67 kDa) from bacterial cultures (Fig. S3, *B–D*). To determine whether ORF150 possesses auto-ubiquitination activity in vitro and identify which E2 enzymes stimulate this activity, we incubated purified ORF150-His with different E2 enzymes in the presence of all the other necessary components. As shown in Fig. 4*A*, ORF150 constituted pronounced auto-ubiquitination when co-incubated with UBE2D1, UBE2D2, UBE2D3, UBE2K, and UBE2L6, as evidenced by specific polyubiquitination bands. These results indicate that ORF150 functions as an E3 ligase. However, the slight spontaneous ubiquitination in the component containing UBE2D3 or UBE2L6 was observed in the absence of ORF150 (Fig. 4*B*).

**Figure 4.**
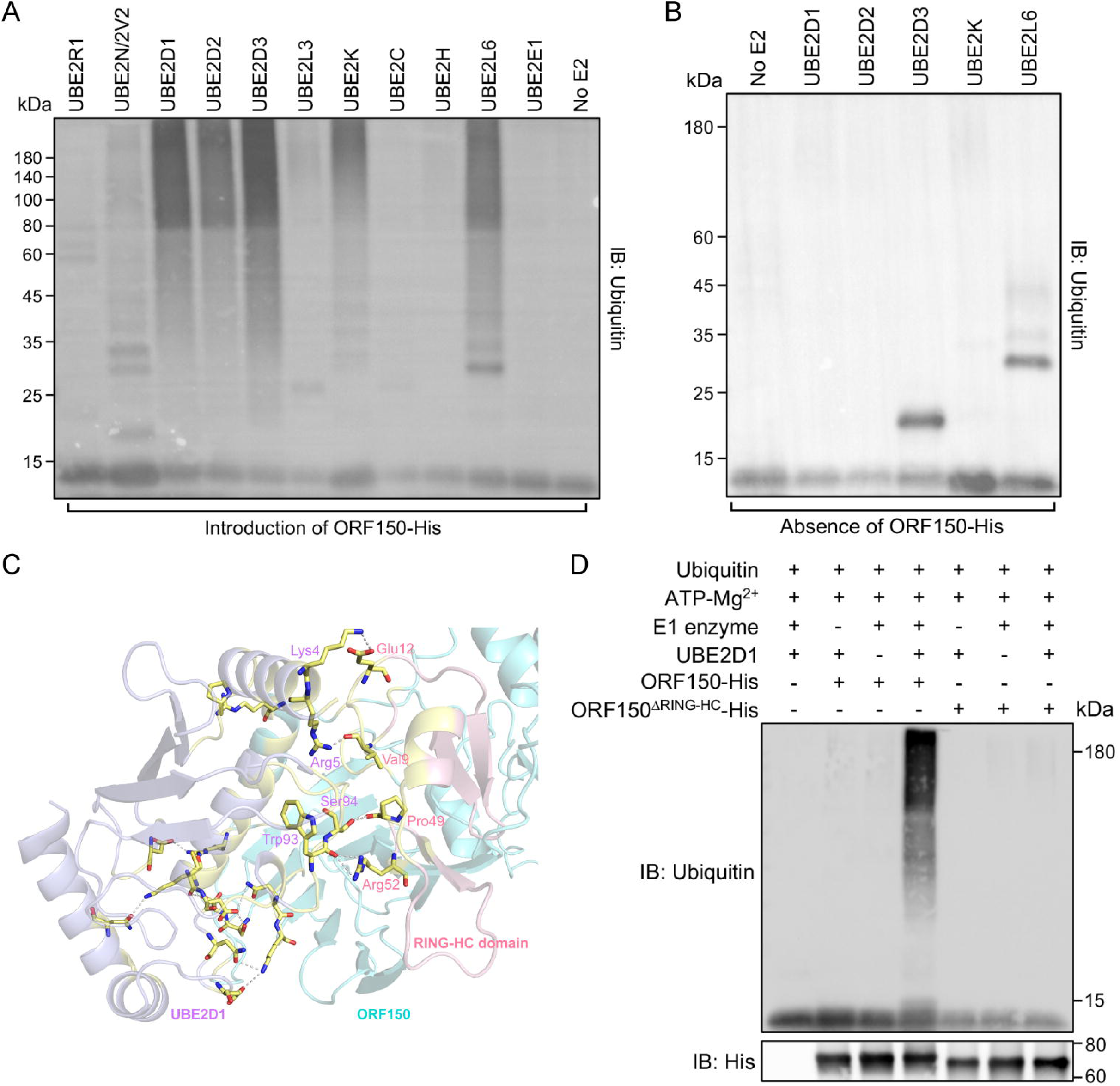
ORF150 functions as an E3 ubiquitin ligase in CyHV-2. *A* and *B*, in vitro ubiquitination assays were performed using anti-ubiquitin antibody under conditions described in the methods. Screening of E2 enzyme activity in the presence of ORF150-His (*A*). Validation of five candidate E2 enzymes (identified in panel *A*) in the absence of ORF150-His (*B*). Negative controls were conducted without E2 enzyme (no E2). *C*, predicted interaction interface between ORF150 (cyan) and UBE2D1 (purple) generated by molecular docking (AlphaFold 3) and visualized in PyMOL. The RING-HC domain of ORF150 was highlighted in pink, and the interaction interface (yellow) displays key binding residues. *D*, in vitro ubiquitination activity of full-length ORF150-His versus its RING-HC domain deletion mutant (ORF150^ΔRING-HC^-His), assessed with ubiquitin, E1 enzyme, UBE2D1, and the other reaction components, followed by immunoblotting with ubiquitin and His-tag antibodies. Negative controls reaction was performed in the absence of E1, E2, or experimental E3 ligases.

Given that UBE2D1 has been extensively validated to stimulate ubiquitination efficiency with a wide range of identified E3 ligases (29), as such, we employed UBE2D1 for subsequent assays. Additionally, computational predictions revealed four stable binding sites between UBE2D1 and ORF150, including interactions within the RING-HC domain (Fig. 4*C*). To clarify the essentiality of the RING-HC domain for ORF150 as an E3 ligase in vitro, we conducted ubiquitination assays using purified full-length ORF150-His and a mutant lacking the RING-HC domain (ORF150^ΔRING-HC^-His, whose size is approximately 62 kDa), as shown in Fig. S3, *B–D*, and then verified by anti-ubiquitin and anti-His antibodies. The results showed that only the reaction containing full-length ORF150 produced a ladder-like smear indicative of high-molecular-weight polyubiquitin chains, while no such pattern was observed in reactions with the mutant deleting RING-HC domain (Fig. 4*D*). These findings confirm that ORF150 functions as an E3 ligase and that the RING-HC domain is essential for its activity.

## Discussion

CyHV-2 as a fish herpesvirus causes severe economic losses in goldfish and gibel carp farming, and to date, no specific control strategies have been developed against CyHV-2, due to its complex infection characteristics (2–7). CyHV-2 induces acute infection in hosts and establishes latent and persistent infections in asymptomatic surviving fishes, which serve as critical sources of infection (5, 6, 30). The underlying mechanisms of CyHV-2 replication and interaction with host remain unclear. Of note, viral proteins directly interact with components of the UPS, which possesses multiple complex mechanisms (31). The UPS is a major cellular mechanism regulating protein degradation and has evolutionarily conserved antiviral defense functions, including the direct degradation of viral components (32, 33). Whereas some viral proteins manipulate the UPS by redirecting E3 ligases to target and degrade host proteins, thereby enhancing viral infection (34, 35). Moreover, foundational studies have demonstrated that many viruses exploit the UPS during their life cycles, including viral entry, gene transcription, assembly, release, and immune evasion (13, 17, 36, 37). However, the potential roles of the UPS in fish herpesvirus infections remain to be confirmed.

In this study, transcriptomics profiling of CyHV-2-infected hosts revealed significant differential regulation of UPS-related genes. Notably, upregulated genes predominated (ubiquitin, proteasome-related components, E1, and E2 enzymes), while more DUBs and E3 ligases was suppressed (Fig. 1, *B–F*). These findings suggest that the UPS is actively involved in CyHV-2 infection; and the virus may establish infection and escape host immune response through suppressing host’s E3 ligases and DUBs (38–40). However, the exact mechanisms is uncertain. Consistent with prior studies, numerous genes associated with the UPS are differentially expressed during SGIV infection (22); during dengue virus infection, the expression level of the E1 enzyme and proteasome subunits were increased (41). Although the UPS plays crucial roles in various viral infections, the underlying mechanisms are different (42–44). For instance, the UPS is required for all stages of human cytomegalovirus (HCMV) infection, as demonstrated by the use of proteasome inhibitors (37). Additionally, HSV-1 capsid undergo cytosolic ubiquitination followed by proteasomal degradation, thereby releasing genomic DNA into the cytoplasm for recognition by DNA sensors (45). Another study reported bortezomib (a proteasome inhibitor) blocks HSV at early steps by halting viral capsid transport to the nucleus and stabilizing host nuclear domain 10 (ND10) structure (46). Herein, the intracellular viral copy numbers and gene transcription levels were significantly reduced with the proteasome inhibitor MG132 treatment, as evidenced by decreased expression of viral proteins observed through IFA (Fig. 2). Notably, based on the temporal characteristics of CyHV-2 gene transcription (47), we analyzed the expression of IE genes, including *ORF54* (encoding a potential E3 ligase with a RING-HC domain) and *ORF121* identified as a highly expressed gene during viral infection (30); E genes, including *ORF62* (encoding potential tegument protein) and *ORF92* (encoding major capsid protein); and L genes, including *ORF33* (encoding a potential DNA packaging terminase subunit 1) and *ORF46* (encoding a potential helicase). The results indicated that MG132 effectively blocks the transcription of viral genes and viral DNA replication. Consistent with findings by Tran and colleagues, who reported that adding proteasome inhibitors at later in HCMV infection prevents further accumulation of viral E and L gene products (42). Therefore, we propose that the host UPS, particularly proteasome, is essential for CyHV-2 infection in vitro.

Notably, to infiltrate the host antiviral defense, viruses employ many strategies, and one of which is manipulating the host’s UPS, where host proteins are redirected for proteasomal degradation (34). This specific process involves E3 ligases that are directly encoded by either the viral or host’s genome (40, 48). This study focuses on the RING-finger family encoded by CyHV-2 genome, revealing that this family is highly conserved among CyHVs. All five members contain a consensus RING-HC domain, a molecular characteristic indicative of E3 ligase activity. These findings suggest that RING-finger proteins are potentially essential for fish herpesviruses. Furthermore, we purified and identified a new E3 ligase, ORF150, encoded by CyHV-2. ORF150 possesses a typical RING-HC domain, which is crucial for its function as demonstrated in Figs. 4*D* and S3, *B*–*D*. Previous studies have demonstrated that E3 ligases exhibit specific preferences for E2 enzymes in proteolytic degradation (49). In contrast, ORF150 appears to display low specificity for E2 enzymes in the ubiquitination assay (Fig. 4*A*), including UBE2D1, UBE2D2, and UBE2K. Notably, slight spontaneous ubiquitination was observed with UBE2D3 or UBE2L6 even in the absence of ORF150. Among these E2 enzymes, UBE2D isoforms serve as ubiquitin donors for a wide range of E3 ligases, involving in critical pathways that influence development and protein turnover, including NF-κB signaling, and DNA methylation, and repair mechanisms (50). Additionally, UBE2K has been implicated in promoting apoptosis and overcoming cell-cycle arrest in response to DNA damage by facilitating the degradation of p53, as well as being associated with multiple neurodegenerative diseases (51–53). The E2 enzymes play pivotal roles in cellular processes and diseases, which are significant for CyHV-2 research. E3 ligases are central determinants of specificity in ubiquitination, which depends on specific E2-E3 and E3-substrate interactions, and the specificity of E2-E3 interaction is rested with only a few amino acid residues (54, 55). As observed in ubiquitination assays, the low specificity of ORF150 for multiple E2 enzymes is a distinctive feature, indicating that the mobilization capacity of ORF150 is more extensive. This property of ORF150 may contribute to the complex characteristics associated with CyHV-2 infection, although further experimental evidence is required to confirm this hypothesis. Similar findings have been reported in studies on WSSV249 (25) and host aspect (56). In addition, the transcription of *ORF150* is significantly upregulated during CyHV-2 infection (Fig. S3*A*). Recent report confirmed that structural variants of *ORF150*, which cover coding sequences of RING-HC domain, exhibit dynamic accumulation during successive passages of CyHV-3 in vitro (57). ORF150 may target NF-κB to inhibit host immune responses, and its deletion attenuates CyHV-3 replication efficiency and virulence in vivo (58). ICP0, a RING-HC type E3 ligase encoded by HSV-1, hijacks the UPS to disrupt cellular pathways regulating host immunity and homeostasis, thereby creating a favorable environment for viral genome replication (59). These observations suggest that evolutionarily conserved ORF150 as an E3 ligase may play a key role in pathogenesis of CyHV-2 infection and its interaction with host, but the specific mechanisms require further investigation.

Overall, we have confirmed that the host UPS is activated during CyHV-2 infection and that the proteasome is essential for viral replication. Additionally, we identified a novel, highly conserved RING-HC domain-containing E3 ligase, ORF150, encoded by CyHV-2, which exhibits a unique low specificity for multiple E2 enzymes. These findings provide the first molecular evidence for the involvement of the host UPS in fish herpesvirus replication and offer new insights into the functions of viral E3 ligases. Our results suggest that targeting the UPS pathway and viral E3 ligases could be promising strategies for developing antiviral interventions against CyHV-2 infection.

## Experimental procedures

### Cell culture, virus propagation, and inhibitor treatment

The *Carassius auratus gibelio* caudal fin (GiCF) cell line and CyHV-2 YC-01 strain (GenBank accession number: MN593216) were isolated and cultured in our laboratory (60, 61). As previous description, CyHV-2 was propagated in GiCF cells cultured in sterile plastic flasks containing 1× Medium 199 (M199; Gibco, USA), supplemented with 2% (v/v) fetal bovine serum (FBS; Gibco) and 1% (v/v) of a 100× antibiotics solution containing 10,000 U/mL penicillin and 10,000 μg/mL streptomycin (Gibco), at 25 ℃. Infected cells were harvested along with the supernatant after complete cytolysis to prepare CyHV-2 stocks and were stored at –150 ℃ (62). For the inhibition assay, GiCF cells were preincubated with DMSO (Sigma-Aldrich, USA) or various concentrations of MG132 (Selleck, USA) for 2 h prior to infection with CyHV-2 at a multiplicity of infection (MOI) of 0.5 for 72 h. Subsequently, whole cell lysates were collected to assess changes in relevant indicators.

### Transcriptomic analysis

To explore the expression pattern of host genes under CyHV-2 infection, de novo RNA sequencing was performed on gibel carps (13–15 cm in length) infected with CyHV-2. Briefly, tissues from both healthy and CyHV-2-infected gibel carps were collected in triplicate, and total RNA was extracted using a TRIzol Reagent (Invitrogen, USA) according to the manufacturer’s protocol. cDNA libraries were constructed using the TruSeq Stranded mRNA Kit (Illumina, USA) and sequenced on the Illumina Hiseq 4000 platform (Illumina) by LC-Bio Co., Ltd. (Hangzhou, China). After removing adaptor sequences, short reads, and low-quality sequences, the clean reads were assembled into transcript sequences using Trinity (63), followed by annotation based on various protein databases as previously described (12). To identify significantly DEGs between the two sample sets, unigene expression levels were normalized to reads per kilobase of transcript per million mapped reads (RPKM), with p-values adjusted for false discover rate (FDR) (64). Unigenes with a fold change (FC) greater than 2 and an FDR less than 0.05 were classified as DEGs (fold change between challenged and unchallenged samples).

### Real-time PCR analysis

For the detection of intracellular gene transcription, total RNA was extracted using a TRIzol reagent (Invitrogen), followed by cDNA synthesis with the PrimeScript™ RT Master Mix (Takara, Japan). For viral copy number quantification, DNA was isolated from whole cell lysates using the TIANamp Genomic DNA Kit (TIANGEN, Beijing, China). Real-time PCR was conducted using the TB Green^®^ Premix Ex Taq™ II (Takara) on a qTOWER^3^ G Real-Time PCR Thermal Cycler (Analytik Jena, Germany), according to the manufacturer’s protocol. Target gene expression levels for intracellular gene transcription was normalized to *β-actin* gene, and the primers were listed in Table S1. The primers for *β-actin* and *ORF121* were used as previously described (47). For viral copy number calculations, a standard curve was generated between copy numbers and cycle threshold (Ct) values based on the DNA polymerase catalytic subunit encoded by *ORF79* of CyHV-2, using primers as previously described (9). The standard curve equation is y = -3.1394x + 38.9, where y represents Ct and x represents the log value of viral copy numbers (R^2^ = 0.9974). All experiments were performed in triplicate.

### Immunofluorescence assay

GiCF cells were cultured on coverslips in 12-well plates and subsequently infected with CyHV-2 (MOI = 1) after pretreatment with either MG132 at 20 µM or DMSO (as a control). At 48 hours post-infection (hpi), the cells were washed with 1× PBS and fixed with 4% paraformaldehyde for 10 min. Non-specific binding sites were blocked using QuickBlock™ Blocking Buffer (Beyotime, Shanghai, China) for 10 min. The cells were then incubated overnight at 4 ℃ with the primary antibody (anti-ORF150, prepared in this laboratory; 1:1000 dilution, as indicated in the figures). Following this, the cells were stained with ABflo^®^ 405-conjugated Goat anti-Rabbit IgG (H+L) (ABclonal, Wuhan, China; 1:300 dilution) for 2 h. After washing with 1× PBS, the cell samples were sequentially counterstained with aggregation-induced emission (AIE) probe (AIE Institute, Guangzhou, China): C_33_H_30_N_4_O_2_^2+^ (for nucleus staining in green) and C_35_H_26_N_3_S_2_ (for membrane staining in red) for 10 min each. Images were captured using Leica SP8 confocal microscope system (Leica, Germany) and analyzed with LAS X Office software (Leica).

### Bioinformatic analysis

The nucleotide or amino acid sequence alignments of the five RING-finger proteins encoded by CyHVs were conducted using Multiple Alignment using Fast Fourier Transform (MAFFT) or Clustal Omega via SnapGene software v4.3.7 (GSL Biotech LLC, USA) or Geneious Prime v2022.0.2 (Biomatters, New Zealand). The conserved domains of the five proteins encoded by CyHV-2 were predicted using the Simple Modular Architecture Research Tool (SMART; https://smart.embl.de accessed on November 20, 2024) (65). Consensus motifs of these proteins were visualized using WebLogo 3 (https://weblogo.threeplusone.com/ accessed on November 25, 2024) (66). Furthermore, phylogenetic trees for the five proteins across CyHVs were constructed using the neighbor-joining method in MEGA v11, with bootstrap values based on 1000 replications (protein IDs are detailed in the figure legends). The potential interaction between ORF150 and UBE2D1 proteins was predicted through molecular docking using AlphaFold 3 (https://alphafoldserver.com/ accessed on December 5, 2024) (67), and the results were analyzed using PyMOL software.

### Plasmids, prokaryotic expression, and protein purification

The coding sequence (CDS) for ORF150 (protein ID: QIV66963.1) encoded by CyHV-2, as well as the mutant ORF150^ΔRING-HC^ (with the RING-HC domain deleted from position 19–153 bp), were synthesized and inserted into the pET-28a(+) vector, incorporating a 6×His tag at the C-terminus (GenScript, Nanjing, China). These plasmids were then used to transform the *E. coli* strain BL21 (DE3) (ABclonal). The transformed cultures were grown overnight at 37 ℃. Subsequently, the cultures were diluted 1:100 in fresh Luria-Bertani (LB) medium supplemented with 40 μg/mL kanamycin (Sangon Biotech, Shanghai, China) and incubated at 37 ℃ to an OD600 of 0.6–0.8. The cells were then induced with 0.4 mM isopropyl-β-D-thiogalactopyranoside (IPTG) (TransGen Biotech, Beijing, China) at 37 ℃ for 6 h.

For the purification of His-tagged proteins, the washed cell pellet was lysed and sonicated in lysis buffer (20 mM Tris-HCl, pH 8.0, 300 mM NaCl, 0.5% NP-40, 10 mM imidazole), followed by clarification through centrifugation at 12,000 *×g* for 30 min. The supernatant containing the His-tagged fusion proteins were subjected to immunoprecipitation using BeyoGold™ His-tag Purification Resin (Beyotime) according to the manufacturer’s protocol, and eluted with elution buffer (20 mM Tris-HCl, pH 8.0, 300 mM NaCl, 250 mM imidazole). The eluted proteins were then dialyzed against a buffer (20 mM Tris-HCl, pH 7.4, 150 mM NaCl, 0–8 mol/L urea) and concentrated using PEG2000 (Sangon Biotech). The un-induced and induced bacterial lysates, as well as the purified proteins, were then verified on 12% SDS-PAGE followed by Coomassie blue staining (Beyotime) and immunoblot (IB) analysis. For western blotting, samples were separated by SDS-PAGE, transferred to a membrane, and probed with a mouse monoclonal antibody against the His-tag (Proteintech, Wuhan, China; 1:5000 dilution for IB) and horseradish peroxidase (HRP)-conjugated goat anti-mouse IgG (H+L) (Epizyme Biotech, Shanghai, China). Protein bands were visualized using a chemiluminescence detection system (Bio-Rad, USA).

### Auto-ubiquitination assay

The purified proteins were used for ubiquitination assays to evaluate their potential E3 ligase activity. Initially, we aimed to identify the E2-conjugating enzymes that significantly promote ubiquitination when bound to ORF150. To this end, we screened 11 types of E2 enzymes using the E2 Select Ubiquitin Conjugation Kit (Yeasen, Shanghai, China) according to the manufacturer’s introduction. The selected E2 enzymes included UBE2R1, UBE2N/2V2, UBE2D1, UBE2D2, UBE2D3, UBE2L3, UBE2K, UBE2C, UBE2H, UBE2L6, and UBE2E1. Then, we employed UBE2D1 to bind with purified ORF150-His and the mutant ORF150^ΔRING-HC^-His to confirm the E3 ligase activity of ORF150 and the necessity of the RING-HC domain. Specifically, in vitro ubiquitination assays were conducted in a buffer containing 10% (v/v) reaction buffer, 10% (v/v) ATP-Mg^2+^, 20% (v/v) ubiquitin, 10% (v/v) E1 enzyme, 5% (v/v) E2 enzymes, and 10% (v/v) purified ORF150-His or ORF150^ΔRING-HC^-His (2 µg). The reactions were incubated at 37 ℃ for 2 h. Each reaction was terminated by adding 5× loading buffer, followed by boiling for 10 min. The samples were then analyzed on 12% SDS-PAGE and subjected to western blotting using rabbit monoclonal antibody against ubiquitin (ABclonal; 1:2000 dilution for IB) or mouse monoclonal antibody against His-tag (Proteintech), followed by incubation with HRP-conjugated goat anti-mouse (Epizyme Biotech) or anti-rabbit (HuaBio, Hangzhou, China; 1:50000 dilution for IB) IgG (H+L).

### Statistical analysis

Data for quantification analyses were shown as mean ± standard deviation (SD). Statistical analyses were conducted using one-way analysis of variance (ANOVA) test via SPSS Statistics v29.0.1.0 (IBM, USA), with significance levels denoted as follows: * *P* < 0.05, ** *P* < 0.01, *** *P* < 0.001, **** *P* < 0.0001, and n.s. indicates no significance (*P* > 0.05). All data are representative of three independent experiments.

## Data availability

Data will be made available on request.

### Supporting information

This article contains supporting information.

### Author contributions

H. W. and J. Y. conceptualization; H. W. and J. Y. methodology; J. Y. and M. L. investigation; H. W. and J. Y. formal analysis; J. Y. visualization; J. Y. and M. L. validation; J. Y. software; H. W., J. Y., and M. L. writing–original draft; H. W., J. Y., and Y. Z. writing–review & editing; H. W. and J. Y. data curation; H. W. and L. L. supervision; L. L. project administration; H. W. and L. L. funding acquisition; H. W. resources.

### Funding and additional information

This work was supported by the National Natural Science Foundation of China [No. 32102842, to H. W.] and the Earmarked Fund for China Agriculture Research System [No. CARS-45-16, to L. Lu.]

## Abbreviations

UPS: ubiquitin-proteasome system
CyHV-2: cyprinid herpesvirus 2
RING: really interesting gene
ORF: open reading frame
KSHV: Kaposi’s sarcoma-associated herpesvirus
SGIV: Singapore grouper iridovirus
HSV-1: herpes simplex virus 1
ICP0: infected cell protein 0
WSSV: white spot syndrome virus
DEGs: differentially expressed genes
psma: 20S proteasome subunit α
psmb: 20S proteasome subunit β
psmc: 26S proteasome ATPase regulatory subunit
psmd: 26S proteasome non-ATPase regulatory subunits
psmf: proteasome inhibitor subunit
psmg: proteasome assembly chaperone
pomp: proteasome maturation protein
DUBs: deubiquitylating enzymes
Usp: ubiquitin-specific protease
uba1: ubiquitin-like modifier-activating enzyme 1
ube2a: ubiquitin-conjugating enzyme E2 A
trim: tripartite motif containing
rnf: RING finger protein
tmem189: transmembrane protein 189
FC: fold change
FDR: false discover rate
RPKM: reads per kilobase of transcript per million mapped reads
DMSO: dimethyl sulfoxide
IE: immediate-early
E: early
L: late
IFA: immunofluorescence assay
AIE: aggregation-induced emission
Cys (C): cysteine
His (H): histidine
CDSs: coding sequences
HCMV: human cytomegalovirus
ND10: nuclear domain 10
GiCF: *Carassius auratus gibelio* caudal fin
FBS: fetal bovine serum
MOI: multiplicity of infection
Ct: cycle threshold
Hpi: hours post-infection
MAFFT: Multiple Alignment using Fast Fourier Transform
SMART: Simple Modular Architecture Research Tool
LB: Luria-Bertani
IPTG: isopropyl-β-D-thiogalactopyranoside
IB: immunoblot
HRP: horseradish peroxidase
SD: standard deviation
ANOVA: analysis of variance.

