## Supplemental Figure S1, S2, and S3; Table S1 for "Investigating E3 ubiquitin ligase activity of ORF150 in fish herpesvirus pathogenesis"

The supporting material list: Figure S1, S2, and S3; Table S1.

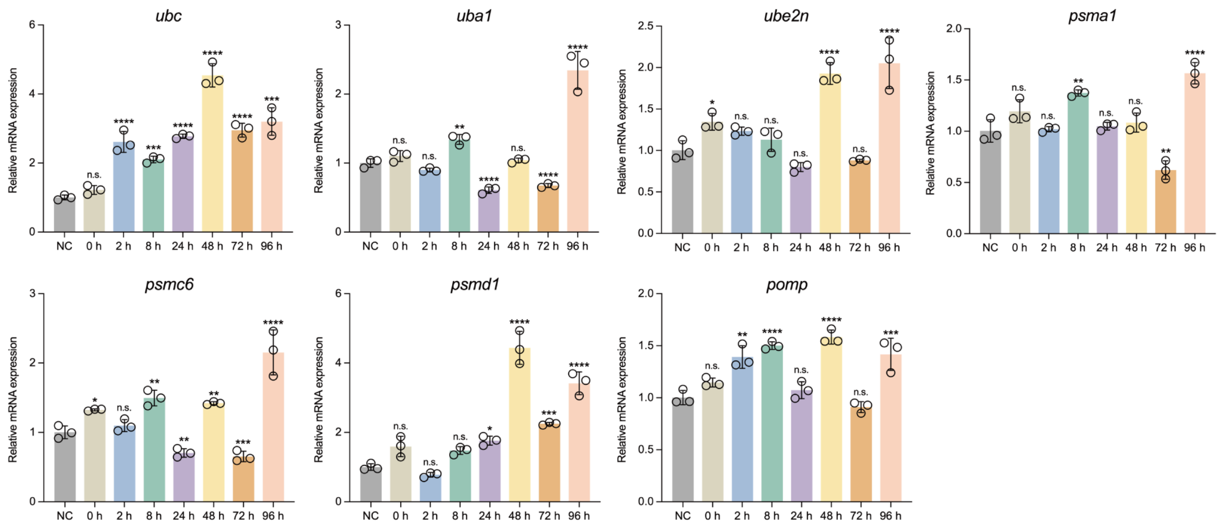


**Figure S1. Ubiquitin-proteasome system (UPS) is activated during cyprinid herpesvirus 2 (CyHV-2) infection in vitro, related to Figure 1.** The changes of transcriptional expression of several UPS-related genes during viral replication in cell model by real-time RT-PCR analysis. Data are representative of three independent experiments. Mean ± SD, statistical analysis was performed using one-way analysis of variance (ANOVA). $^{*}$ $P < 0.05$, $^{**}$ $P < 0.01$, $^{***}$ $P < 0.001$, $^{****}$ $P < 0.0001$, and n.s. indicates no significance ($P > 0.05$).

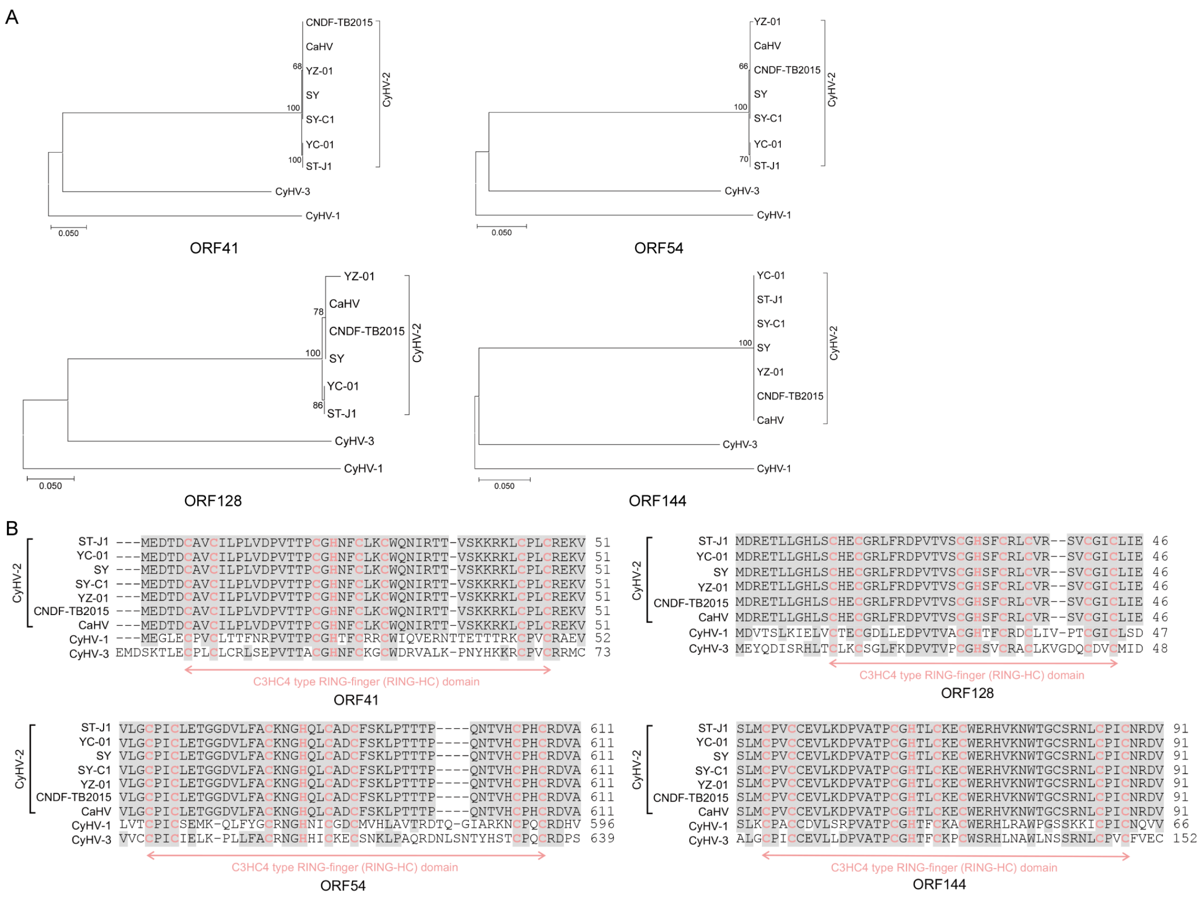


**Figure S2. CyHV-2 encodes highly conserved RING-finger proteins, related to Figure 3.** *A*, phylogenetic trees of open reading frame (ORF)41, 54, 128, and 144 (including homologs) among CyHVs were constructed using the neighbor-joining method in MEGA v11 with bootstrap values of 1000 replications. *B*, the multiple sequence alignments of ORF41, 54, 128, and 144 proteins among CyHVs (complete amino acid sequences are not shown), and the consensus residues were covered by grey shade. The RING-HC domains are marked in pink. In *A* and *B*, protein IDs are following: for ORF41 (including homologs), CyHV-2 ST-J1, YP_007003862.1; SY-C1, AKC01991.1; SY, AMB21612.1; YC-01, QIV66859.1; YZ-01, QAU54767.1; CNDF-TB2015, QIM55193.1; *Carassius auratus* herpesvirus (CaHV), APB92892.1; CyHV-1, YP_007003707.1; CyHV-3, YP_001096076.1; for ORF54 (including homologs), CyHV-2 ST-J1, YP_007003875.1; SY-C1, AKC02003.1; SY, AMB21625.1; YC-01, QIV66872.1; YZ-01, QAU54779.1; CNDF-TB2015, QIM55210.1; CaHV, APD51570.1; CyHV-1, YP_007003720.1; CyHV-3, YP_001096089.1; for ORF128 (including homologs), CyHV-2 ST-J1, YP_007003954.1; SY, AMB21704.1; YC-01,

QIV66951.1; YZ-01, QAU54857.1; CNDF-TB2015, QIM55310.1; CaHV, APD51590.1; CyHV-1, YP_007003783.1; CyHV-3, YP_001096163.1; for ORF144 (including homologs), CyHV-2 ST-J1, YP_007003959.1; SY-C1, AKC02082.1; SY, AMB21709.1; YC-01, QIV66956.1; YZ-01, QAU54862.1; CNDF-TB2015, QIM55323.1; CaHV, APB92987.1; CyHV-1, YP_007003797.1; CyHV-3, YP_001096179.1.

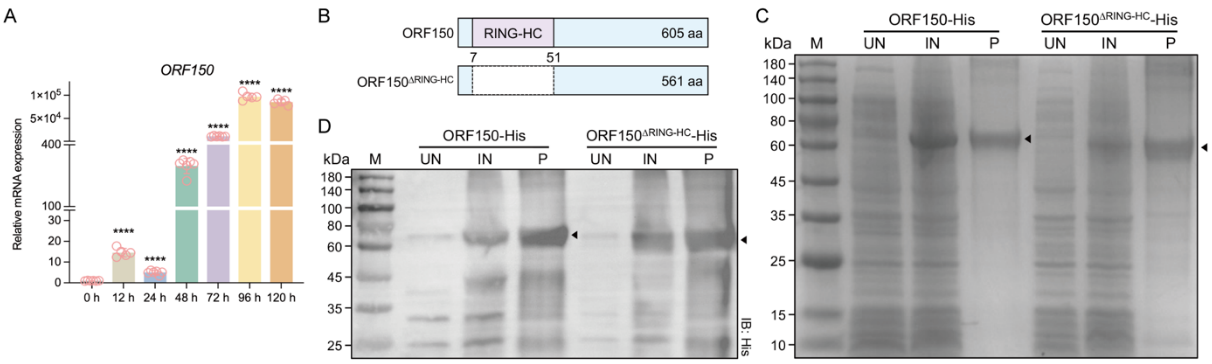


**Figure S3. The expression pattern of *ORF150*, and purification of ORF150 and its domain-deleted mutant, related to Figure 4.** *A*, the changes of intracellular transcriptional expression of *ORF150* during viral replication by real-time RT-PCR analysis. Data are representative of three independent experiments. Mean ± SD, statistical analysis was performed using one-way ANOVA. **** $P < 0.0001$. *B*, Schematic representation of full-length ORF150 protein and mutant constructs. *C* and *D*, the un-induced (UN) and induced (IN) bacteria solution, and purified proteins (P) were verified on 12% SDS-PAGE by Coomassie blue staining (*C*) and western blot analysis (*D*) with His-tag antibody. M, protein marker.

**Table S1. The primers used in real-time PCR assays.**

| Genes | GenBank accession number | Primer sequences (5’ to 3’) |
|---|---|---|
| *ubc** | XM_052592323.1 | F: AGCGACACCATTGAGAAC<br>R: AATGTGCGACCATCTTCC |
| *uba1** | XM_052594119.1 | F: GGTGTAGAGATAGCCAAGAAT<br>R: CGCCAGTGTAGGAGGTTA |
| *ube2n** | XM_052582744.1 | F: CGCTACTTCCATGTGGTCATC<br>R: CGCTTAGTAATGCCTGGATTGA |
| *psma1** | XM_052544517.1 | F: TCGGAACCAGTATGACAATGAC<br>R: ACAGCGTGAGAGCGTGAT |
| *psmc6** | XM_052586426.1 | F: TTGTCGGTGAGGTGCTGAA<br>R: ATCGTGAGAGTCGTCATATCCA |
| *psmd1** | XM_052539168.1 | F: AGACGAGGAGAAGGAGAAGGA<br>R: GCGGACACTGGCTCTACAA |
| *pomp** | XM_052588485.1 | F: GCTGCTACCGAGTCATCCAT<br>R: GAATCTGTCTAGCCGCTCTGTA |
| *ORF33* | CyHV-2 YC-01 strain MN593216.1 | F: GGCGTTCAGTATAGCACAACAA<br>R: GCATCCGACACCACCTTGA |
| *ORF46* | | F: GAAGGTTCCGAGGAGGTGAT<br>R: TCAGCATCGCAGTCCAGAG |
| *ORF54* | | F: GACATACGCCATCAACAAGGAT<br>R: CTCGCTCGCCACATCATCT |
| *ORF62* | | F: TCTATCAAGTGCTGTCCAACCT<br>R: GAATGTAACCGTCGTCGTATCC |
| *ORF92* | | F: GAGCCACCTTCCTGTTCCA<br>R: GACGAGCAGCGAGTAGTGA |
| *ORF150* | | F: TCCCAACATGATCAGGTGCC<br>R: GAACACGCGCTATCTGAGGA |

* Abbreviations: ubc, polyubiquitin-C; uba1, ubiquitin-like modifier-activating enzyme 1; ube2n, ubiquitin-conjugating enzyme E2 N; psma1, 20S proteasome subunit α 1; psmc6, 26S proteasome ATPase regulatory subunit 6; psmd1, 26S proteasome non-ATPase regulatory subunit 1; pomp, proteasome maturation protein.
